# Hyperspectral Oblique Plane Microscopy Enables Spontaneous, Label-Free Imaging of Biological Dynamic Processes in Live Animals

**DOI:** 10.1101/2023.03.15.532804

**Authors:** Ke Guo, Konstantinos Kalyviotis, Periklis Pantazis, Christopher J Rowlands

## Abstract

Spontaneous Raman imaging has emerged as powerful label-free technique for investigating the molecular composition of biological specimens. Although Raman imaging can facilitate understanding of complex biological phenomena *in vivo*, current imaging modalities are limited in speed and sample compatibility. Here, we introduce a single-objective light-sheet microscope, named ***λ***-OPM, which records Raman images on a timescale of minutes to milliseconds. To demonstrate its function, we use ***λ***-OPM to map and identify micro-plastic particles based on their Raman spectral characteristics. In live zebrafish embryos, we show that ***λ***-OPM can capture wound dynamics at five-minute intervals, revealing rapid changes in cellular and extracellular matrix composition in the wounded region. Finally, we use ***λ***-OPM to obtain Raman scattering maps of a zebrafish embryo’s beating heart at an effective 28 frames per second, recording compositional changes at different points in the cardiac cycle.

## 1 Main

The Raman effect occurs when a photon couples to a molecular vibrational transition as it scatters from an object. This scattered photon has a different wavelength to the incident photon; plotting the spectrum of the scattered photons renders peaks, the shape and distribution of which are highly characteristic of the sample and its biochemical composition. This selectivity can be used to identify cancer cells[1], investigate protein aggregation in Parkinson’s Disease[2] and Alzheimer’s Disease[3], study the molecular dynamics of apoptosis[4] and even probe the extraordinary adhesive qualities of the mussel byssus[5].

Despite the advantages of this technique for non-invasive and label-free materials characterization, such as in medical diagnosis, archaeology or forensic investigation, a major shortcoming is speed. The spontaneous Raman cross-section of biomaterials is typically small, requiring integration times of 0.1 to 10 s per measurement. Raman mapping, in which a Raman spectrum is gathered for every location in a microscope’s field-of-view, can easily take hours and is therefore insensitive to faster biological processes, such as muscle activity, neural signalling, gamete fusion, flagellar motion, platelet aggregation or a multitude of other high-speed processes besides. Part of the reason for the low speed of spontaneous Raman scattering detection is the inefficient use of non-Raman-scattered photons; the vast majority of incident photons pass through the focus unchanged, providing no appreciable signal but incurring a substantial risk of photodamage. Light-sheet microscopy (LSM), in which the sample is illuminated from the side by a thin ”sheet” of light and signal is recorded from all points along it, can make use of these photons[6]. Unlike in conventional Raman imaging when the sample is illuminated by a single focussed spot, light-sheet photons which pass through the periphery of the sample may yet undergo Raman scattering from locations further along the light-sheet’s direction of propagation. Nevertheless, LSM suffers from difficulties with sample mounting. Conventionally, the light-sheet is projected from a secondary microscope objective mounted orthogonally to the primary objective. The limited room between the two objectives sets significant constraints to the sample size and geometry, focusing or moving the sample can be difficult, and the commercial solutions lack flexibility in adaptation or modification. Because Hyper-spectral LSMs have been limited to dual-objective designs until now, they have seen little uptake amongst the imaging community, and have not yet fully exploited the speed advantages that this imaging configuration promises[7–10].

To overcome these shortcomings we present a single-objective epi-illumination hyperspectral light sheet microscope, *λ*-OPM, which possesses the throughput advantages of a hyperspectral LSM while resolving the sample-mounting complications. It is based on an oblique-plane microscope design[11] which uses a single microscope objective to both project the light-sheet and capture the scattered light. This makes it compatible with most established sample formats, as well as commercial accessories such as stage-top incubators. We demonstrate the full sensitivity and spectral characterization capabilities of *λ*-OPM by taking 3D-resolved Raman spectra from dispersed mixtures of microplastics and classifying them by their spectral fingerprint. Furthermore, to showcase the ability of *λ*-OPM to record dynamic processes, we monitor a wound developing in a zebrafish embryo at unprecedented five-minute intervals. Finally, we observe the heartbeat in a live zebrafish embryo using Raman contrast at an equivalent frame rate of 28 frames per second.

## 2 Results

### 2.1 Design of a single-objective epi-illumination hyperspectral light-sheet microscope

Biological phenomena occur on a wide range of timescales, and many cellular processes are too fast to be imaged using existing Raman microscopes. To address the need for fast, label-free Raman imaging of dynamic biological phenomena, we set out to design a microscope that can capture spontaneous Raman maps in minutes, or even seconds, without requiring exotic sample mounting geometries. Our design is based on an oblique plane microscope format (**Fig. 1**); a laser beam is projected through the sample at a tilted angle, and emission light (such as fluorescence, Raman scattering or Brillouin scattering) is captured by the same objective. An image of this tilted plane is formed using a second microscope objective, and a third objective is then used to compensate for the tilt, by placing the whole tilted plane within its depth of focus. For details of the components used, see **Online Methods**.

**Fig. 1.**
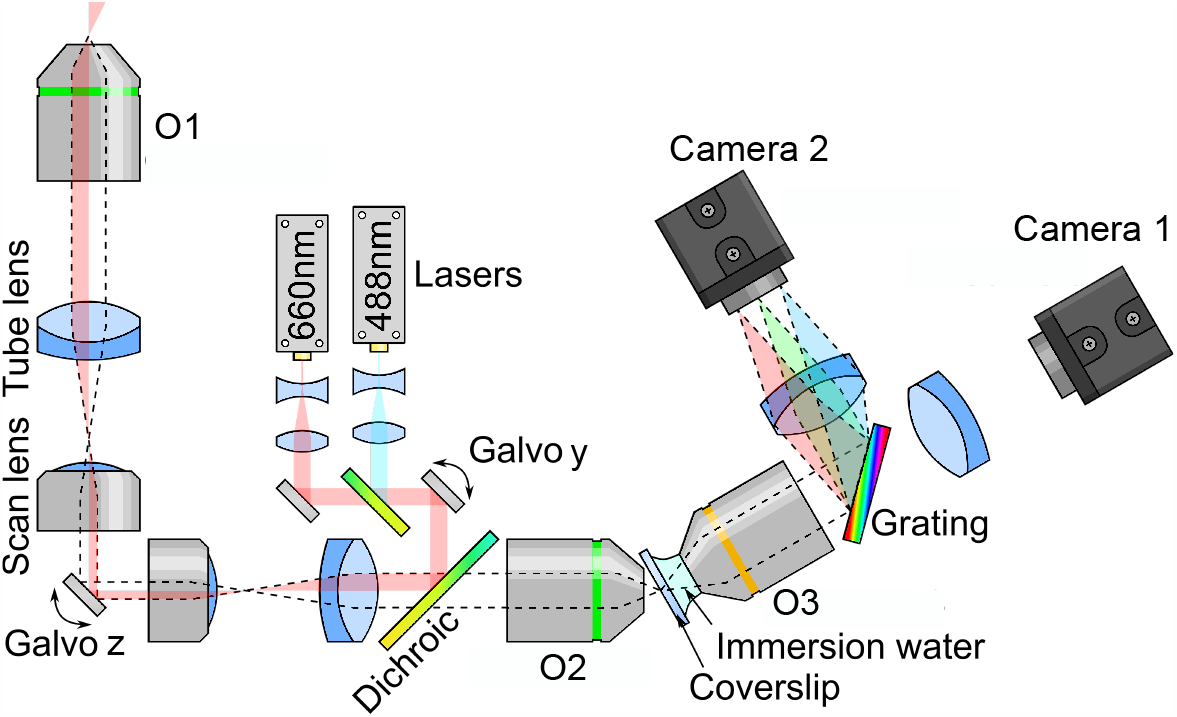
Optical diagram of the single-objective epi-illumination hyperspectral light-sheet microscope, *λ*-OPM. Light from one of two lasers (488 nm or 660 nm) is expanded to ∼ 0.5 mm diameter before being brought to a focus at the image plane of a microscope. The focus can be scanned anywhere in the image plane using two orthogonal galvanometric mirrors (Galvo y and Galvo z), which are made conjugate to each other using a 4f lens relay made from a commercial scan lens and tube lens. This image is relayed to the sample with a tube lens and microscope objective (O1). Because the laser illumination passes close to the edge of the system aperture, the sample is illuminated by an oblique, weakly-focused beam. The light emitted or scattered from the sample is then imaged back through the optical system, through the dichroic mirror and into a second microscope objective (O2) where it forms a virtual image with a magnification equal to the refractive index of the sample (∼ 1.33). A third objective (O3) then images this tilted plane onto a camera via a tube lens. For hyperspectral imaging a grating is placed in the optical path to disperse the light from the illuminated line by wavelength, recording an optical spectrum for each illuminated point. The grating can be replaced with another filter cube containing a dichroic mirror and two bandpass filters for more conventional two-colour imaging. For details on the components used, see **Online Methods**.

*λ*-OPM incorporates a grating into the infinity path of the third objective, turning it into an imaging spectrometer, with an entrance slit defined by the image of the obliquely-projected laser beam. In conventional imaging mode the system has a field of view of 500 μm × 400 μm with a measured resolution of 650 nm laterally and 3400 nm axially full-width half-maximum (FWHM) (**Supplementary Fig. 4**) within the centre ∼ 250 μm × 250 μm of the field of view; image quality is degraded slightly for objects located deeper within the sample. In hyperspectral imaging mode, the Y axis is used as a spectral axis, providing a spectral range of approximately ∼ 200 nm, corresponding to Raman shifts of up to ∼ 3500 cm^*−*1^ with our 660 nm excitation laser. The center wavelength can be changed either by manually adjusting the grating tilt or by moving the laser beam position with Galvo y. The vertical resolution is approximately 10-20 μm (FWHM) from the centre to the left and right edges of the image as determined by the laser beam width (see **Supplementary Section 2**).

### 2.2 λ-OPM maps and identifies arbitrary microplastics in minutes

A major virtue of Raman microspectrometry over conventional bright-field or fluorescence microscopy is the ability to uniquely identify the major biochemical constituents of almost any sample without the need for labelling. To demonstrate this functionality, we measured the Raman spectra of a selection of different polymer micro-particles and thereby determined their composition.

We used a convolutional neural network (CNN) [12, 13] to classify the different polymers. It was trained on measured spectra of the three polymers, agarose and the spectrum of the glass-bottomed dish as detailed in **Online Methods**. In **Fig. 2(b)** we show a map of 198 stage locations in the y direction, with brightness and colour indicating the signal strength and the type of material respectively: Polystyrene in red, Poly(methyl methacrylate) (PMMA) in green, Polyamide / Nylon-6 (PA6) in blue, and agarose or dish in grey. Compared to a white light scattering-contrast image of the same area in **Fig. 2(c)**, the map captures not only the location and shape of the particles in the focal plane with high contrast, but also the composition of each. The Raman spectra of the three marked pixels in **Fig. 2(d)** show clear correspondence with previously-reported reference spectra [14, 15], verifying the material classifications. Overall, we demonstrate that *λ*-OPM enables fast, label-free accurate identification and classification of various polymers, offering a powerful and precise means of material analysis and characterisation.

**Fig. 2.**
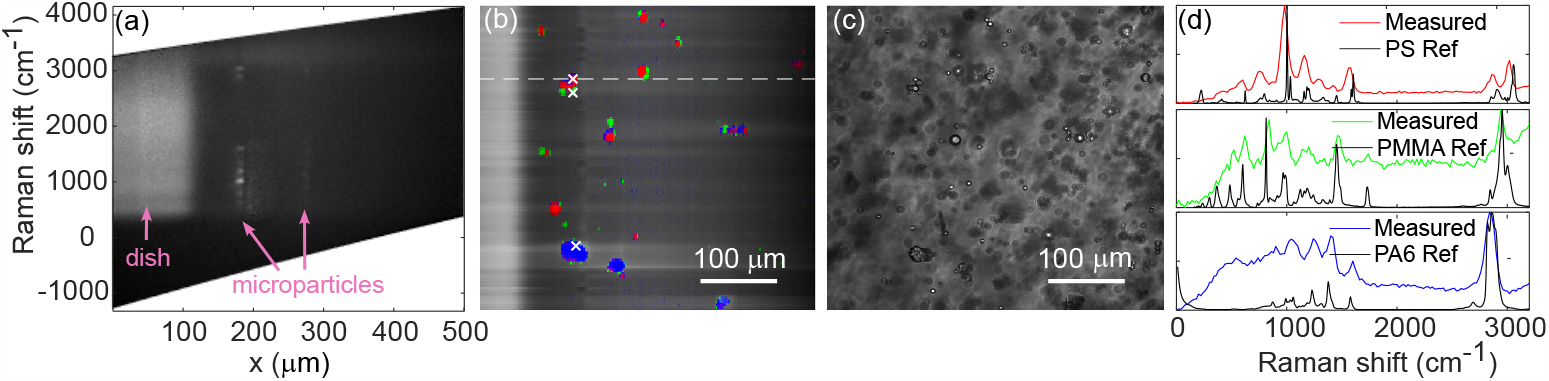
*λ*-OPM performs Raman mapping and identification of microplastics in less than two minutes. (**a**) A hyperspectral image showing two different spectra of polystyrene (PS) and poly(methyl methacrylate) (PMMA). (**b**) A false colour Raman scattering image constructed from 198 scan locations in the y direction, with the type of material at each position identified using a convolutional neural network (CNN). The brightness of the image is scaled with a gradient in the x direction to compensate for the laser intensity reduction as it passed through the sample. Red: PS, green: PMMA, blue: polyamide / Nylon-6 (PA6), grey: agarose or dish. The data was binned every 10 pixels in the x direction. The measured line highlighted by the dashes corresponds to the dataset shown in (**a**). (**c**) A (Rayleigh) scattering image of the same sample area illuminated by a white Light-Emitting Diode (LED) from above the sample stage. (**d**) Measured Raman spectra at the three marked pixels in (**b**), showing good correspondence to the reference spectra of the corresponding polymers identified by the CNN. Note the horizontal axis in units of wavenumbers, or waves per centimeter. Exposure time: 0.5 s per line, laser power: 800 mW.

### 2.3 λ-OPM enables rapid *in vivo* Raman imaging of wound dynamics in zebrafish embryos

Apart from quickly characterising and identifying static materials, the real advantages of *λ*-OPM are label-free imaging of dynamic, *in vivo* processes. To showcase this capability, we measured the 2D-resolved Raman spectra of live, wounded zebrafish embryos every five minutes, during a 1-2 hour period after wounding. A single incision was made in the dorsal myotome opposite to the anal pore using a pair of forceps, as described in **Online Methods** (**Fig. 3(a)**). In **Fig. 3(b)** we show exemplar Raman spectra both at the wound and in the muscle next to the wound (serving as a control) at two different timepoints: one early post-wounding (27 min) and one at a later stage (61 min). The laser beam profile was deconvolved out of the measured spectra to improve spectral resolution, and the autofluorescence signal was estimated from the measured spectra and subtracted to increase peak contrast (for details see **Supplementary Section 6**).

**Fig. 3.**
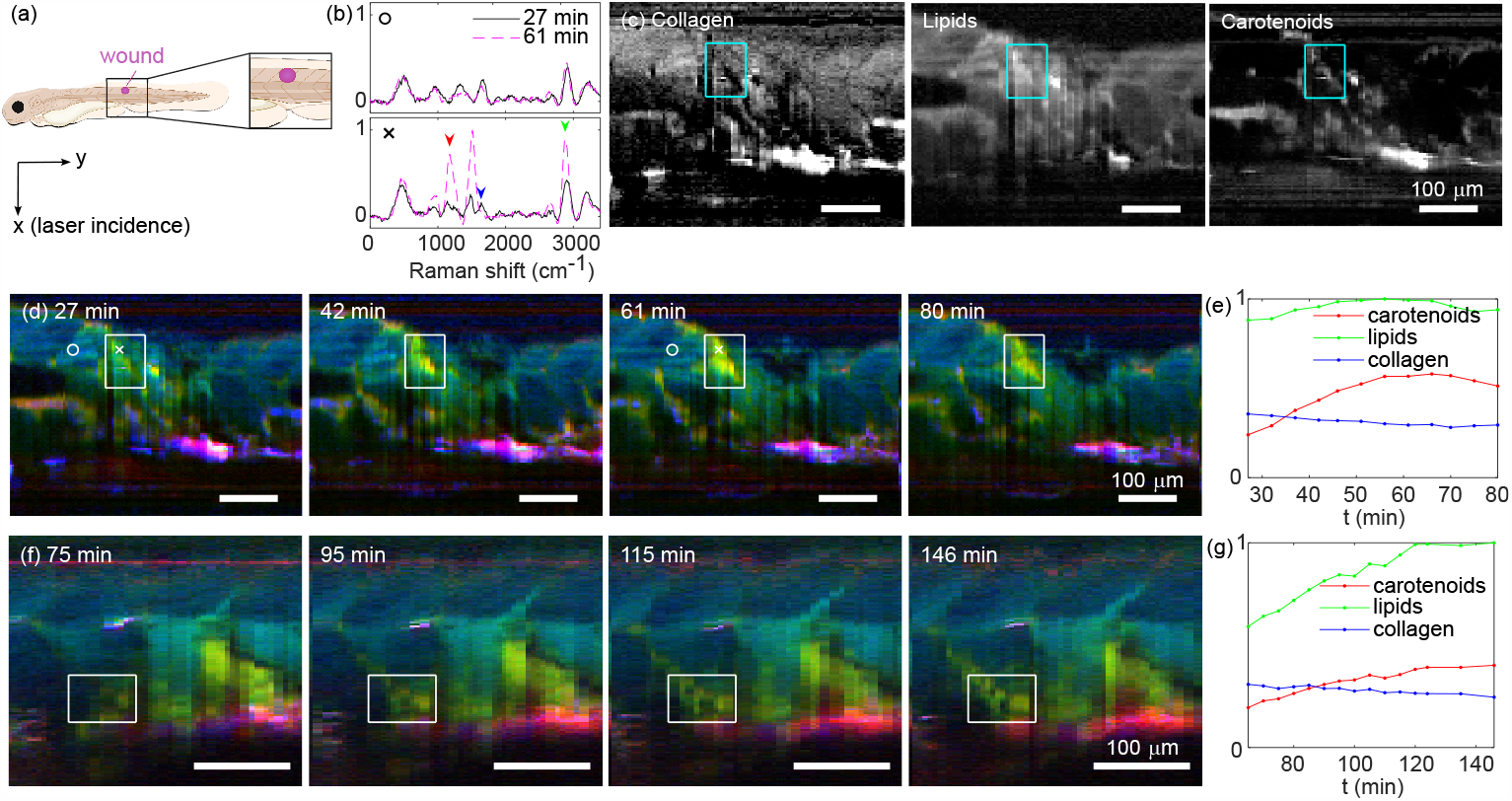
*In vivo* Raman imaging of wound development in zebrafish embryos allows the tracking of important biomolecules, including lipids, collagen and carotenoids. (**a**) An illustration of the investigated zebrafish embryo showing the estimated image area and wound location. (**b**) Example Raman spectra (normalized, with laser beam size deconvolved and fluorescence signal subtracted [23]) obtained at different times post injury. (**c**) Distribution of collagen, lipids and carotenoids determined by averaging the peak intensity (±40 cm^*−*1^) at 1640 cm^*−*1^, 2880 cm^*−*1^ and 1160 cm^*−*1^ respectively. The arrows in (**b**) mark the position of the peaks. (**d,f**) False colour images of the wound areas from two zebrafish embryos at different times after the wounds were made. The red, green and blue channels indicate the distribution of carotenoids, lipids and collagen. The first image (27 min) in (**d**) combines the three images in (**c**). The white circles and crosses mark the positions and times corresponding to the spectra in (**b**). (**e,g**) Normalized intensity of the carotenoids, lipids and collagen peaks averaged over the wound regimes within the squares in (**d**) and (**f**) as functions of time after the wounds were made. Laser power: 600 mW, exposure time: 2 s per frame. Note that (**d**) is taken with approximately 1.5 times the field of view of (**f**).

The (control) muscle spectra do not appear to change significantly over time, with peaks near 960 cm^*−*1^, 1330 cm^*−*1^ and 1670 cm^*−*1^ characteristic of collagen [16]. The peak near 2920 cm^*−*1^ likely corresponds to proteins (2930 cm^*−*1^[17]) or lipids (2880 cm^*−*1^[18]), and the peaks near 3240 cm^*−*1^ are most likely due to water in the tissue [19]. In contrast, the wound exhibits significantly different spectral components, with weaker collagen peaks and two extra peaks near 1160 cm^*−*1^ and 1500 cm^*−*1^. The 1160 cm^*−*1^ peak is characteristic of carotenoid spectra [20], while the 1500 cm^*−*1^ peak can be interpreted as a combination of peaks at 1525 cm^*−*1^ from carotenoids and 1450 cm^*−*1^ from lipids. Both peaks grew significantly higher as the wound development progressed. The peaks near 2920 cm^*−*1^ also increased with a shift to 2880 cm^*−*1^, suggesting an increase in the lipid concentration at the wound site[18].

To map spatiotemporal changes in the relative concentration of carotenoids, collagen and lipids near the wound area, we averaged over the peaks (± 40 cm^*−*1^) at 1160 cm^*−*1^, 1640 cm^*−*1^ and 2880 cm^*−*1^ respectively. In **Fig. 3(c)** we present the distribution of the three components, which clearly demarcate the wound site from the adjacent undamaged tissue. Equivalent images can be combined into a single false colour image, to visualize compositional changes over time; **Fig. 3(d)** and **(f)** show representative Raman images taken from two zebrafish embryos at four different timepoints as the wound development progressed. The average intensities of the three components in the wound regime enclosed by the white rectangles are plotted in **Fig. 3(e,g)**. The wound site was characterised in both embryos mainly by an increase in the carotenoid signal, indicating accumulation of blood. Furthermore, the emergence of the wounds was marked by an increase in lipid concentration accompanied by a decrease in that of collagen. This increase in lipid concentration could be attributed to free fatty acids that have been released as part of early-stage wound signalling, as previously reported [21, 22].

With hyperspectral contrast, many factors can be explored to shed light on the mechanisms of wound healing, making *λ*-OPM a powerful new tool for studying the dynamics of such processes non-destructively, *in vivo* and without the need for labelling.

### 2.4 λ-OPM performs Raman mapping of the developing zebrafish beating heart at “video rate”

To showcase the full capabilities of *λ*-OPM in recording very fast processes, we imaged the beating heart in live zebrafish embryos at an effective 28.6 frames per second. To carry this out, a conventional (i.e. non-hyperspectral) video of the heart beating was first taken as a reference, with the laser scanning in y at 200 Hz to form a light sheet, and using Camera 1 with 35 ms exposure time. Afterwards, the laser was positioned near the centre of the image and the grating was inserted for hyperspectral imaging. The hyperspectral images were taken continuously at the same frame rate and exposure time, while every 35 s the zebrafish was translated in the y direction in 5 μm steps using the microscope stage, resulting in approximately 1000 frames per y step. The cardiac motion did not change significantly over the whole measurement time of less than one hour, allowing us to construct a 2D hyperspectral video of the heart by combining the images at the same phase of the *cardiac cycle*, or sequence of events during which the atrium and ventricle sequentially fill with blood and then contract, pumping the blood round the body. The cardiac phase of each hyperspectral image was obtained by cross-correlating the hyperspectral images with the pre-acquired video as detailed in **Online Methods**. For every cardiac phase, 40 images were averaged, reducing shot noise and random read noise by a factor of approximately six. To minimise the influence of the auto-fluorescence on the Raman signal, we averaged over the Raman spectra at 1100-1200 cm^*−*1^ (auto-fluorescence, red channel); the fluorescence subtracted Raman spectra at 2800-2900 cm^*−*1^ (lipids or proteins, green channel) and 3100-3300 cm^*−*1^ (water, blue channel) respectively. In **Fig. 4** we show a sequence of false-colour images at 17 consecutive timepoints in the cardiac cycle (a full cycle takes approximately 14 frames). A video containing the sequence of the heart beating can be found in **Supplementary Video 1**.

**Fig. 4.**
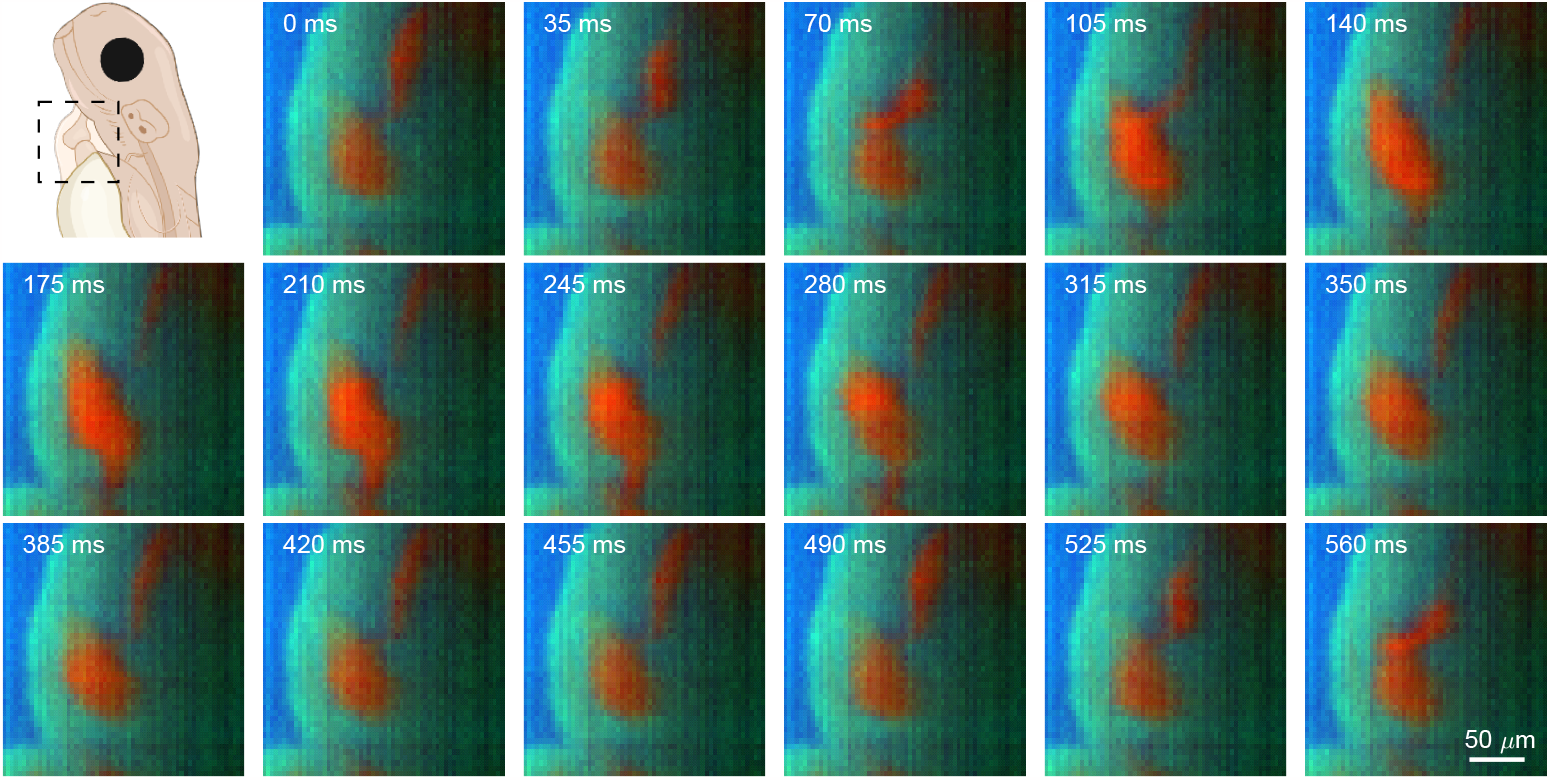
Cardiac synchronization permits *in vivo* Raman imaging of a beating heart in a zebrafish embryo. Reconstructed false colour images of a zebrafish embryo heart at different cardiac phases over slightly longer than one heart beat cycle. The green, blue, and red channels indicate the distribution (normalized) of lipids/proteins, water and auto-fluorescence respectively. Laser power: 600 mW, exposure time: 35 ms per frame.

Overall, the unique design of the *λ*-OPM enables high-speed Raman imaging of the beating heart of a zebrafish embryo, producing comprehensive 2D hyperspectral maps of the cardiac cycle. These capabilities pave the way for spontaneous, rapid and label-free imaging of millisecond-scale biological processes *in vivo*, such as those that occur during embryo development.

## 3 Discussion

*λ*-OPM was designed from the start to achieve unprecedented hyperspectral volumetric imaging performance, in terms of speed, throughput and sensitivity, without compromising the familiarity, flexibility, ease of use and compatibility with thirdparty components that a commercial microscope frame provides. That is not to say that this is the only means to achieve high-speed Raman-contrast images; Coherent Raman Scattering (CRS) microscopy and Surface-Enhanced Raman Scattering (SERS) microscopy both exhibit improved pixel throughput by enhancing the Raman cross-section. Nevertheless, CRS is difficult to implement and expensive, typically requiring synchronized picosecond laser sources and complex detection schemes, and does not typically capture the full spectral content [24]. On the other hand, SERS does capture the full Raman spectrum, but the need for nanoprobes or a SERS substrate means that measurements are arguably not label-free, and may even be limited to measurements of phenomena which occur at the sample surface [25]. As such, spontaneous Raman imaging has a clear role to play in many fields, and *λ*-OPM is able to broaden the range of accessible phenomena to those that occur on the timescale of minutes and seconds.

*λ*-OPM achieves its performance through careful component choice; a 0.93 theoretical collection numerical aperture, low-noise cameras and use of high-performance anti-reflection coatings help maintain image brightness despite the grating dispersion and significant number of optical surfaces present in the system. It is this throughput and sensitivity which gives rise to the high imaging rates as low as effectively 35 ms per line *in vivo*. The design permits the user to change from two-channel to hyperspectral imaging without needing to realign. This capability was achieved by creating a filter cube insert containing a grating on a custom kinematic mount. For two-channel imaging the filter cube can be replaced by one containing a dichroic and two emission filters. Other key design features are the ability to rapidly scan the excitation beam using a galvanometric mirror (for use when rapidly capturing hyperspectral depth maps), and a custom water-immersion chamber for the tilted OPM objective. This water chamber is more complex to implement than a solid immersion lens[26], but offers greater flexibility, larger field of view and the opportunity to use lenses with a broader range of design wavelengths. Depending on which objective lens is chosen, this can result in significant cost savings as well. Further construction details for the grating mount and water-immersion chamber can be found in **Supplementary Section 1**.

In terms of utility, the ability to reliably demultiplex three microplastic samples without the need for labelling shows the power of spontaneous Raman microspectrometry; distinguishing similar molecules requires analysis of the whole Raman spectrum, not just isolated bands. Because of the low Raman cross-section though, gathering data at speed is extremely difficult; capturing high-spatial-resolution Raman images within minutes or even a few hours is typically impossible. *λ*-OPM’s ability to do so is therefore unique. Given the information-rich content of each spectrum it is expected that scaling to larger numbers of microplastics would be readily achievable. With the rising concern over the prevalence of microplastics in the natural environment [27, 28] and the tissues of plants and animals as well as humans [29, 30], instruments like *λ*-OPM can find considerable utility in this burgeoning field.

Previous studies of wound healing have highlighted the speed of the process, with molecular signalling starting as soon as a few minutes after injury [21, 22]. *λ*-OPM can quantify multiple biomolecules (including lipids, carotenoids, collagen and more) simultaneously on this timescale. The two representative embryos we showcase here exhibited subtly different wound progressions in time, which can be attributed to many factors, such as the intrinsic differences between animals, the inevitable differences in the manual wounding operation, and the imaging location relative to the centre of the wound. With the hyperspectral contrast of *λ*-OPM, many factors can be explored to shed light on the mechanisms of wounding and eventually wound healing.

*λ*-OPM operates without labelling by fluorophores or other probes, which is particularly important when investigating complex processes, since parameters such as molecular diffusion and adsorption coefficients[31], kinetics and thermodynamics of protein / ligand interactions[32], solubility and rates of aggregation[33], and even the morphology of neurotoxic amyloid fibrils[34] can all be subtly affected by the fusion of a fluorescent protein or small molecule to a protein of interest. The data themselves show clear biochemical differences compared to adjacent tissue: infiltration of blood and an increase in the concentration of lipids, attributed in this case to signalling by free fatty acids. Tracking these parameters during wound treatment has potential to help develop new treatments for traumatic injury, free from experimental artifacts caused by labelling.

The ability to use Raman contrast to probe biological phenomena occurring on the timescale of milliseconds is perhaps the most ground-breaking achievement of *λ*-OPM. While the approach taken relies strongly on the periodic nature of the cardiac cycle, it is nevertheless flexible enough to accommodate changes in heart rate while providing label-free spontaneous Raman contrast images with a 28 fps “video rate” frame rate. Zebrafish have been used as models for cardiogenesis[35], coronary artery disease[36] and arrhythmia[37], thus being able to study cardiac processes with molecular specificity is a unique new capability for understanding the heart. In the future, Raman mapping of the developing or regenerating beating heart could be used to monitor the intricate processes of cardiac morphogenesis and regeneration, eventually revealing key biochemical changes that could be important for translational science.

While *λ*-OPM is flexible and is compatible with almost all conventional microscopy samples, there are of course limitations. Like all light-sheet systems there is an increased sensitivity to sample scattering compared to confocal and multiphoton microscopy; when performing hyperspectral imaging this scattering can start to degrade the spectral resolution as well (since the spectrum is convolved with the width of the excitation profile). In practice, the effect is manageable and deconvolution can be used to recover even fine details in biological Raman spectra. As case in point, we have successfully imaged almost 400 μm into a live zebrafish embryo, confirming that this sensitivity does not preclude practical biological Raman imaging.

In summary, we have recorded label-free spontaneous Raman maps of several dynamic phenomena, using the first ever single-objective hyperspectral light sheet microscope, *λ*-OPM. As a light sheet technique, it addresses common shortcomings in hyperspectral imaging, such as high photon loss and low speed, while minimizing out-of-focus photodamage and background. Moreover, it is free from the sample mounting complications of conventional light sheet systems. As a label-free technique it requires no sample staining, does not suffer from photobleaching to any reasonable degree, and is compatible with long-term imaging of a wide range of different samples. Label-free recording of uniquely-identifiable Raman spectra was demonstrated by successfully identifying individual microplastic particles in a mixture of three (all within two minutes), and the speed and biological utility highlighted by mapping two different phenomena with data throughput previously deemed unachievable by spontaneous Raman microspectrometry. These were recording a sequence of 500 μm × 350 μm spontaneous Raman maps of wounded zebrafish embryos at five-minute intervals, and recording a beating heart at video-rate *in vivo*.

## 4 Methods

### 4.1 Instrument construction

For reasons of user familiarity and ease of use, *λ*-OPM was constructed as an add-on to a commercial microscope frame (Olympus IX73). A laser beam (LaserQuantum Gem 660 nm 1 W, spatially filtered to achieve a pure TEM_00_ mode profile) is reflected by a dichroic beamsplitter (Semrock Di03-R660) and focused through part of the imaging path to a spot near the edge of the back focal plane of a water immersion objective (O1, 25 × NA=1.1, Nikon MRD77220 mounted in a custom nosepiece), producing a weakly-focused thin line in the sample, tilted at an angle of 38° to the flat surface of the outermost lens element in O1. Scattered light or fluorescence emission from the line is collected by the same objective, through the dichroic beamsplitter and a longpass filter (Semrock LP02-664RU), and ultimately imaged onto the remote focusing plane, with an overall magnification of approximately 1.33×. This overall magnification must be equal to the refractive index of the sample, in accordance with the remote focusing principle [38]. Matching the refractive index is achieved by splitting the penultimate lens into two physical lenses (Thor Labs TTL200MP and AC508-750-A-ML), and tuning their separation to change their effective overall focal length (and thus the magnification of the system). The light then passes into O2 (air immersion, 20× NA=0.7, Olympus UCPLFLN20X) to form a theoretically-perfect remotely-focussed image. The back aperture of O2 is slightly smaller than the image of the back aperture of O1 and a misalignment between them can reduce the resolution and introduce aberrations. Therefore, O2 is mounted on a three-axis translation stage (Newport M-561D-XYZ with two SM-13 micrometers for XY positioning and a DS-4F differential micrometer for Z focusing) for fine position adjustment.

The remote focusing plane lies on the outer surface of a coverslip, which functions as a wall of a custom-made water chamber (see also **Supplementary Section 1**). The image is refracted through the coverslip and water into a third objective (O3, 25 × NA=1.1, Nikon MRD77220) mounted inside the water chamber. The optical axis of O3 is tilted by 38° so that the remote plane is normal to its optical axis, and the translation stage supporting O2 is used to match the field of view of O2 and O3. The refraction by the water chamber negates the NA loss due to the mismatch of the light cones of O2 and O3 caused by the tilt [39]. The line illuminated by the laser is finally imaged horizontally (in the x direction) on to a camera with a total magnification of 33. For the spectral resolution, a horizontal grating (ThorLabs GR25-0310, 300 lp/mm, 8.4 ° blaze angle, tilted by about 5°) together with a longpass filter (Semrock LP02-664RU) is inserted using a custom insert (see **Supplementary Section 1**) in a commercial kinematic fluorescence filter cube (ThorLabs DFM1/M) between O3 and the tube lens before a second camera. This grating diffracts the m = -1 order image in the y direction as a function of wavelength, creating a hyperspectral image with as many as 2560 spectra captured simultaneously (i.e. one spectrum per column in the image). Subsequent to the grating, two different tube lenses are used for different measurements. For the microplastics and part of the zebrafish embryo measurements (Fig. 2 and Fig. 3(f-g)) we used a standard 200 mm tube lens (ThorLabs TTL200-A), resulting in a magnification of 33× in the x direction. For most measurements with zebrafish embryos (Fig. 3(b-e) and Fig. 4), we used a compound lens consisting of four achromatic doublets (ThorLabs AC508-400-B) combined to yield an effective focal length of approximately 100 mm. The magnification is reduced by a factor of about 2 in both directions to improve the SNR against the read noise of the camera.

### 4.2 Imaging Capacity

*λ*-OPM can capture volumetric hyperspectral data by scanning the position of either the sample or the laser beam. Two galvanometric mirrors (Galvo y and Galvo z), both placed conjugate to the pupil of O1, can be used to control the position of the laser line. Galvo y scans the laser line in the y direction. When dithering back and forth in high speed, the laser line forms a light sheet and a whole plane in x-y can be imaged, as in a conventional OPM. Galvo z scans the laser line (or sheet) in z and de-scans the image while keeping tilt of the plane constant [40]. The sample is mounted on a motorized scanning stage (Thorlabs MLS203), which can be used to scan the sample in both x and y.

The field of view of the OPM is 500 μm × 400 μm. The maximum spectral range of a hyperspectral image is approximately 200 nm with the centre of the spectrum determined by the vertical location of the laser beam, i.e. scanning Galvo y changes the spectral range. For Raman mapping, we keep Galvo y fixed.

The theoretical detection NA of the OPM image is approximately 0.93, with the theoretical limiting aperture being the back aperture of O2. By imaging a sparse distribution of 210 nm diameter fluorescence beads (Tetraspeck microspheres, 0.2 μm blue/green/orange/dark red) in agarose gel, we found the actual resolution of the OPM to be approximately 650 nm × 650 nm × 3400 nm (FWHM) in the middle of the image; the resolution degrades towards the edge of the image due to stronger aberrations as shown in **Supplementary Fig. 4**. When doing hyperspectral imaging, the resolution in the y direction is determined by the size of the laser beam, i.e. 8 to 20 μm (FWHM, from centre to edges). This also determines the spectral resolution to be between 13 and 40 nm. When imaging large objects, spectral resolution can be improved by deconvolving the laser profile. In the case of the zebrafish wound measurements, a Gaussian function with FWHM of 18 *μ*m was used as an estimate of the laser profile. It was deconvolved from the measured image using the Matlab deconvlucy function. When imaging sparsely-distributed objects that are smaller than the laser beam, the spectral resolution is not limited by the beam size but by the vertical size of the object. However, the position of the object relative to the laser beam can introduce a shift of the spectra. Deconvolving the object image improves both the spectral resolution and accuracy. In addition, as a general challenge for light sheet microscopy, scattering of the excitation light in the sample can reduce the contrast of the image as well as the spatial and spectral resolution. Where necessary, the spectral resolution could be further enhanced by adding a slit on the remote image plane between O2 and O3, at a cost of photon loss.

### 4.3 Spectral calibration

In a hyperspectral image, the horizontal position (x) corresponds to the position along the imaged line, while the vertical position (y) is approximately a linear function of the wavelength and the vertical position of the laser beam, i.e. the setting of Galvo y. The tilt of the grating further causes a vertical shift of the image that is proportional to x. Therefore, given the setting of Galvo y and the grating constant (300 lp/mm), the corresponding wavelength in the final hyperspectral image can be expressed as a function of x and y with two unknown parameters, i.e. the grating tilt and a prefactor describing the x dependent vertical shift of the image.

A coarse calibration was first conducted by measuring the transmitted fluorescence of an aqueous quantum dot solution (CdSeS/ZnS alloyed 665 nm from Sigma-Aldrich) through the longpass filter (Semrock LP02-664RU). An estimate of the two calibration parameters was obtained by matching the transmission edge to the cut-off wavelength of the filter. Then we measured Raman spectra of PS and PMMA. By comparing a few measured spectra to online data [14, 15] with the estimated parameters as starting values, we adjusted the parameters until an optimal match of the peak positions was achieved.

As the tilt of the grating can change subtly after removing and re-inserting the cube, for maximal spectral accuracy, re-calibration is necessary every time the cube is re-inserted. To facilitate re-calibration for zebrafish measurements, we measured a standard Raman spectrum of zebrafish embryo muscle without moving the grating cube using the calibration parameters obtained from the PS and PMMA spectra. When re-calibration is required, the water and collagen peaks of a new measured muscle spectrum are compared with the corresponding peaks of the standard muscle spectrum to obtain the new calibration parameters. The same set of parameters is used for all data measured without moving the grating cube.

### 4.4 Zebrafish embryo heart beat measurements and data processing

A sequence of Raman images showing a full heart beat cycle(s) was obtained from over 45000 hyperspectral images taken continuously with 35 ms exposure time (28.6 fps). During the image acquisition, the sample was moved every 35 s to each position in y using the translation stage, i.e. approximately 1000 frames per y position. As the camera and the translation stage were not synchronized, we only use the middle 800 frames to avoid including images at a “wrong” y position. All images were first preprocessed, including background subtraction, rotation to compensate for camera tilt, and cropping.

In order to locate the frames at the same phase in the heart beat cycles, we used Camera 1 to record a 100-frame video of the heart beat with the same frame rate as a reference, assuming that the heart beat pattern did not change significantly over the whole measurement time (less than 1 h). This was done before the scan and can also be done afterwards. The video images are resized to match the pixel and y step size of the scan and then cropped to the same regime. A few exemplar pre-processed video images can be found in **Supplementary Fig. 6**.

For each y position (*N*_*y*_ positions in total), we took the corresponding 800 hyper-spectral images and summed over the spectral range (-40 to 3400 cm^*−*1^), yielding sequences of rows which we combined into *N*_*y*_ images with vertical direction showing the time variation. We would then process these images independently by comparing them to the corresponding rows of the video from Camera 1 as following. First, in order to reduce the influence of the noise, we further removed the columns without significant time variation, i.e. only columns with standard deviation larger than 80 % of the average standard deviation are left. For each time point of the heart beat cycle, we took the row of interest from 11 consecutive frames of the video before and after this time point (slightly shorter than a full cycle) to form an image, which also showed the time variation in the vertical direction. We cross-correlated this image with the corresponding time variation scan image. The peaks of the cross-correlation indicated which subsections of the scan image has a similar time variation pattern to the video image. They were expected to be in the centre in the horizontal direction. The vertical position of the peaks provided the numbers of the scan frames at the same heart beat phase. We selected the highest 40 peaks of the cross-correlation and summed the corresponding frames to get an average hyperspectral image for the corresponding time point and y position. Repeating for a sequence of time points and combining results for different y positions, we obtained the hyperspectral information of a full heart beat cycle which would then be processed in the same manner as the wound data.

The hyperspectral images were then processed into spectra. After subtracting the fluorescence signal (see **Supplementary Section 6**), the different peak intensities were calculated from the spectral data for every pixel at every time point to form the final Raman image sequence. **Supplementary Fig. 7** shows a comparison between a few preprocessed video images and the corresponding final Raman images.

### 4.5 Convolutional neural network for microplastics classification

The CNN used for classifying the Raman spectra of microplastics follows the template in Ref. [13] using one layer with filters = 64 and kernel size = 3. The training and test data were measured from 3 samples, each with one type of the polymer particles in agarose, along with one sample containing pure agarose for the spectra of agarose and any residual signal from the glass-bottomed dish. Hyperspectral images were taken at different positions of both galvo mirrors and then converted to spectra with 20 cm^*−*1^ resolution and with binning of 10 pixels in the x direction. The Raman and fluorescence spectra of agarose and the dish were obtained from the corresponding positions of the hyperspectral images measured from the pure agarose sample. The Raman spectra of PS, PMMA and PA6 were programmatically selected based on the the number of peaks in the spectra. In detail, we smoothed each vertical line of the image to reduce noise (after rotation) and search for the peaks (MATLAB smoothdata and findpeaks) in two separate regimes, i.e. 700-1850 cm^*−*1^ and 2700-3200 cm^*−*1^. When at least 3/4/3 (for PS/PMMA/PA6) peaks in the first regime and at least 2/1/1 (for PS/PMMA/PA6) peaks in the second regime were found, the corresponding spectrum was calculated as training data.

The whole data set contains about 15000 spectra (approximately 3000 for each class). The data were randomly shuffled and pre-processed, which included removing the linear trend (using scipy.signal.detrend) of the spectra, subtracting the mean, normalizing to the maximum and finally cropping to the range of 560 cm^*−*1^ to 3360 cm^*−*1^. 80 % of the data were used for training and 20 % for testing. The final model converged after 200 epoches and reached an accuracy of over 98 % after 500 epoches. **Supplementary Fig. 5** shows the convergence of the training process.

For each spectrum, the CNN output a set of 5 scores corresponding to the 5 materials, which we used to classify the Raman spectra at different locations. Using the 5 scores as weights, the total scattering intensity was converted to the RGB channels of the image, with the PS channel in red, PMMA in green, PA6 in blue, and agarose or dish (sum of the two divided by 3) for all three channels (grey).

### 4.6 Sample preparation

#### 4.6.1 Polymer particles in agarose

The PS (6-10 μm from Sigma-Aldrich), PMMA (6-10 μm from Goodfellow), and PA6 (5-50 μm from Goodfellow) particles were pre-mixed with a ratio of about 1:1:1 using a vortex mixer. Approximately 0.004 g of the polymer mix was mixed into 1 mL hot 2 wt% agarose gel, and immediately dropped into a confocal dish to create a pad. The samples used for CNN training were prepared in the same way, by separately adding a measured quantity of each polymer particle composition into 2 wt% agarose gel, with slightly higher concentration to optimise particle density.

#### 4.6.2 Zebrafish husbandry

Experiments involving zebrafish were conducted in accordance with UK Home Office requirements (Animals Scientific Procedures Act 1986, project license P219D3ABD). All experiments were conducted up to 5 days post-fertilization (dpf). At 24 hours post fertilisation (hpf), PTU (Merck) was added to the E3 medium in order to reduce pigmentation.

#### 4.6.3 Zebrafish myotome wound experiments

Wild-type (WT/AB) embryos at 3 dpf were anaesthetised in 0.2 mg/ml MS222 (Merck) and mounted on 1.2 % w/v/ low-melting point agarose (Merck). The embryos were wounded by making a single incision using a pair of forceps (Dumont No. 5) in the dorsal myotome opposite to the anal pore. After wounding, the embryos were transferred to egg medium (E3) to allow for recovery. Prior to imaging, embryos were anaesthetised and subsequently mounted on 0.7 % w/v low-melting point agarose (Merck) in 20 mm glass-bottom Petri dishes (VWR). A hair loop was used to position the embryos such that the wound was close to the glass surface of the Petri dish. After gelation, each dish was supplemented with ∼1-2 ml of E3 containing 0.2 mg/ml MS222.

## Supporting information

Supplementary video 1: Heart beating

## 5 Data availability

A CAD model of the instrument along with analysis software, characterization data and the datasets used for each figure can be found at Zenodo using the following links.

https://doi.org/10.5281/zenodo.8368477 (Microplatics experiment)

https://doi.org/10.5281/zenodo.10009113 (Zebrafish heart)

https://doi.org/10.5281/zenodo.8380484 (Zebrafish wound)

https://doi.org/10.5281/zenodo.10045008 (CAD model and PSF)

## 6 Acknowledgements

CJR acknowledges funding from EPSRC (EP/S016538/1, EP/X017842/1 and EP/W024969/1), BBSRC (BB/T011947/1 and BB/X004716/1), Wellcome Trust (212490/Z/18/Z), Cancer Research UK (29694 and EDDPMA-May22\100059), the Royal Society (RGS \R2 \212305 and IES \R2 \222231), the Chan Zuckerberg Initiative (2020-225443 and 2020-225707) and the Imperial College Excellence Fund for Frontier Research. PP acknowledges funding from BBSRC (BB/T017929/1), MRC (MR/X019837/1) and the Royal Society Wolfson Merit Award (WRM\FT\180010).

## 7 Author contributions statement

Research was conceived by CJR and PP. *λ*-OPM was designed and constructed by CJR and KG, animal handling was performed by KK, experiments were carried out by KG with the assistance of KK. Data analysis was performed by KG with guidance from CJR, PP and KK. All authors contributed to the manuscript.

