## Supplementary Information for "Hyperspectral Oblique Plane Microscopy Enables Spontaneous, Label-Free Imaging of Biological Dynamic Processes in Live Animals"

### 1 System design

#### 1.1 Water chamber fabrication

In any OPM design, O3 must be tilted with respect to O2, which creates obvious difficulties in capturing the rays propagating at high angles to the optical axis of O3. Traditionally this has been achieved with a high-NA-air-immersion objective (which cannot achieve the NA necessary to capture all the rays emitted by O2), a high-NA water immersion objective with a custom-made water chamber [1] or more recently a dedicated solid-immersion zero-working-distance OPM objective[2, 3]. Due to the cost and inflexibility of zero-working-distance lenses, we use a water chamber solution. An existing water-immersion objective (Nikon MRD77220, 20×, 1.1NA) was fitted with a custom-machined water chamber (see Supplementary Fig. 1(A),(B)) which sat over the lens, and could be positioned accurately using a differential micrometer. O-ring seals ensured the water chamber did not leak, and the outermost optical surfaces consisted of glass coverslips cut to shape and glued onto the chamber housing. A separate mounting clamp held the water chamber tightly to the objective body. A translation stage attached between the clamp and the water chamber was used to adjust the position of O3 so that it focused on the outer surface of the coverslip to minimise spherical aberration.

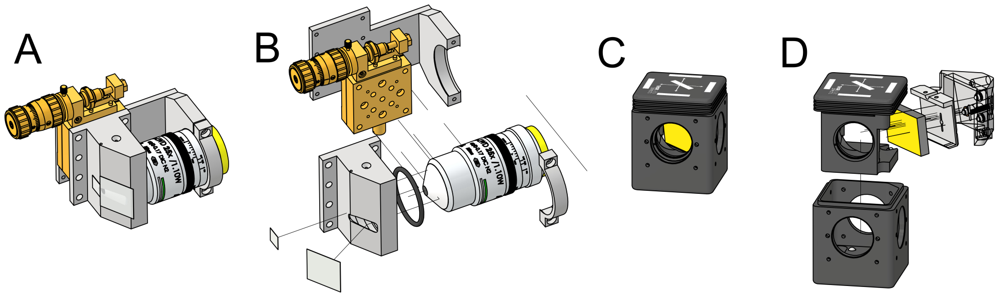


**Supplementary Figure 1 Custom parts.** (A) Assembled water chamber assembled onto a microscope objective. The focal position of the coverslip can be tuned using a differential micrometer. (B) Exploded diagram of (A). (C) Modified fluorescence filter cube containing a grating mounted in a custom kinematic mount. (D) Exploded version of (C), highlighting the custom-machined modified insert, grating mount, three-point kinematic adjusters and retaining spring.

#### 1.2 Grating insert fabrication

λ-OPM is designed to rapidly and easily convert between a conventional OPM design and the hyperspectral imaging configuration. A custom insert for a commercial kinematic fluorescence filter cube (ThorLabs DFM1/M magnetic filter cube) was created by modifying the existing DFM1T1 cube insert. One half of the insert was replacing with a custom-machined kinematic grating mount, and the existing half was milled to make room (see Supplementary Fig. 1(C),(D)). The insert held a grating oriented such that the zero order reflection passes outside the system aperture. The first order diffraction propagates along the optical axis, through the tube lens and images onto the camera. The three-point kinematic adjusters and retaining spring allow for small change of the grating tilt angle and therefore the spectral range detected by the camera.

A separate filter cube, containing a 532 nm dichroic mirror (Semrock DI03-R532-T1-25X36) and two band-pass filters (Semrock FF01-515/30-25 and FF01-582/64-25), could be used instead of the grating cube, allowing easy conversion between the hyperspectral imaging mode and conventional red and green filter channels.

#### 2 Laser beam profile

The laser beam profile was characterized by taking the fluorescence image of an aqueous solution of quantum dots (Sigma-Aldrich CdSeS/ZnS alloyed 665 nm, diluted) using Camera 1, assuming the laser beam to be circularly symmetric. The following figure shows an example beam profile. After binning the image by 10 in the horizontal direction, we calculate the FWHM of each vertical cut, which is 10 μm in the centre and less than 18 μm on the left and right side of the image.

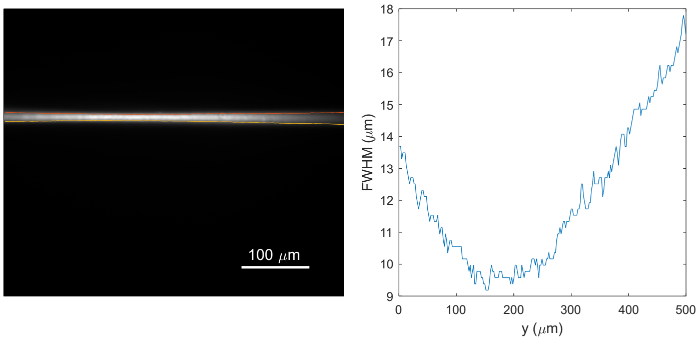


**Supplementary Figure 2 Laser beam profile.** (Left) A fluorescent image of the laser beam measured from an aqueous solution of quantum dots. The red and yellow lines indicate the positions of the half maximum. (Right) FWHM of the laser beam as a function of the horizontal position.

### 3 Background scattering

The optics near the intermediate image plane introduced some strong fluorescence and scattering spots in the hyperspectral image frame. While these could be eliminated by subtracting a blank image, imprecise subtraction (due to sample effects and laser fluctuations) and photon shot noise occasionally resulted in artefacts (see Supplementary Fig. 3) or spurious intensity fluctuation at the corresponding x position in the final map, e.g. in Fig. 2(a).

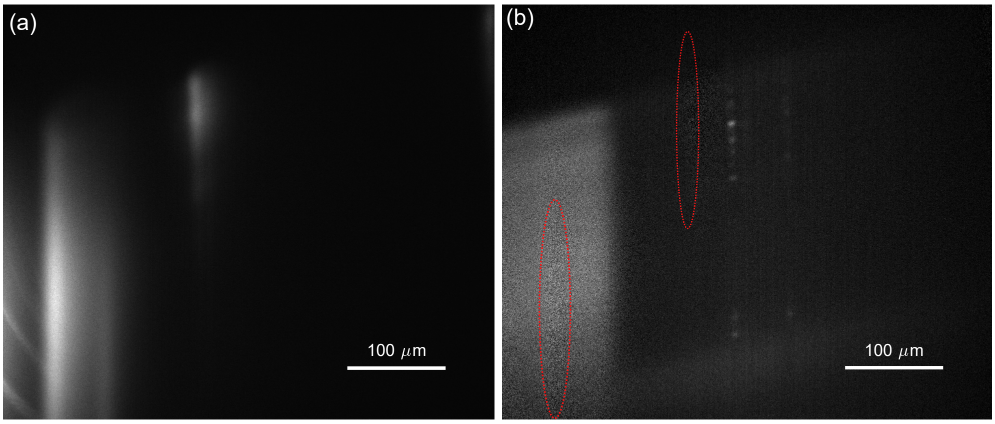


**Supplementary Figure 3 Background scattering.** (a) An image of the background scattering from intermediate optics measured with a beam block placed between the commercial microscope frame and the customized parts. (b) An hyperspsectral image showing artefacts (red circles) due to errors in background subtraction.

### 4 Point spread function (PSF)

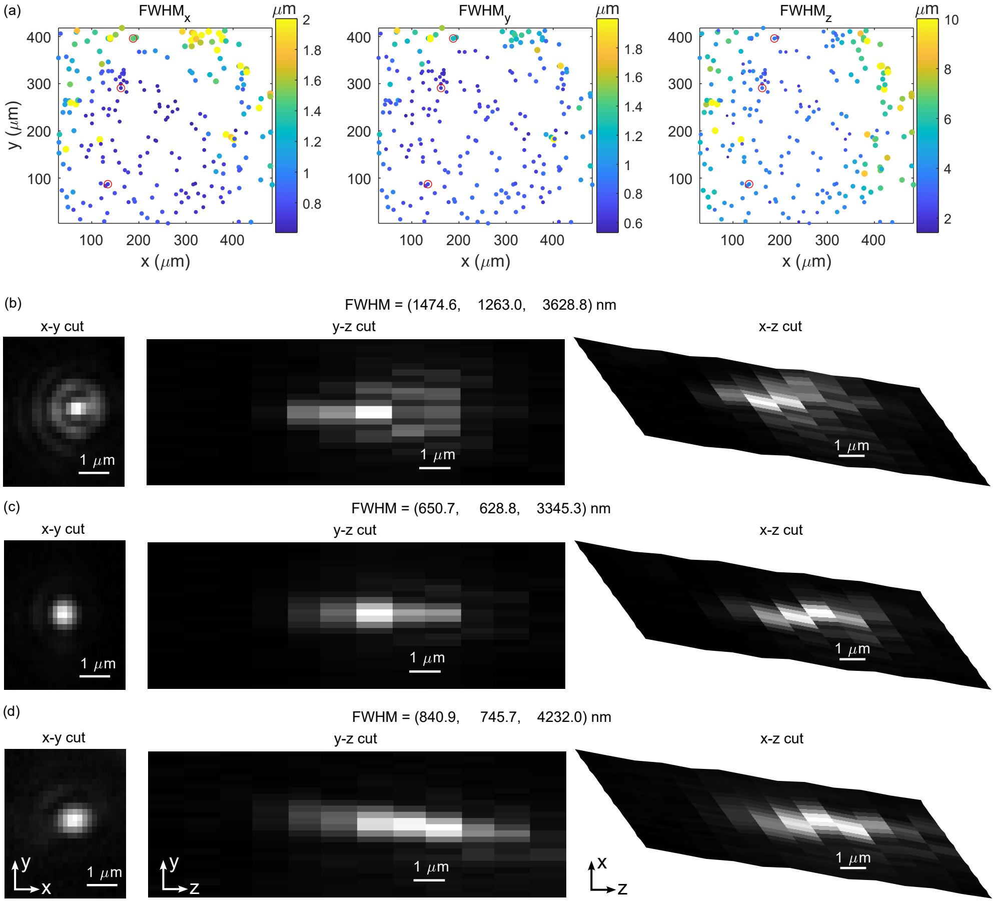


**Supplementary Figure 4 Point spread function.** (a) Distribution of the FWHM of the 3D PSF at different positions of the image. The resolution is evenly distributed in the middle 250 μm × 250 μm area and increases at the edges of the image. The values are obtained by fitting each the 3D images of 224 fluorescent beads with 3D Gaussian functions (least square method). (b-d) Sections in the middle of the images of the 3 beads marked by red circles in (a) ordered from the top to the bottom of the field of view. The larger FWHM results from stronger aberrations near the edge of the field of view. Laser power: 5 mW, exposure time: 2 s.

The PSF was measured from the images of 210 nm fluorescent beads (Tetraspeck microspheres, 0.2 μm blue/green/orange/dark red) embedded in agarose. To prepare the sample, the bead suspension was diluted in water by 20 times, mixed with 2 % Agarose solution (warm), and finally on a 35mm glass-bottomed dish to form a pad. The sample was excited using a blue laser (Coherent Obis 488nm LX 150mW) with Galvo y dithered to form a lightsheet. The beamsplitter and filter set was replaced by a beamsplitter and filter set for matching wavelengths (FF497-Di01 and FF01-565/133-25). Images were taken at a few Galvo z settings with 800 nm steps to form a 3D image (after correcting for the skew). To obtain the FWHM of the PSF, the 3D image of each bead was fit individually by a 3D Gaussian function using least square method. The FWHMs of each bead image in the x,y and z directions were calculated from the fitting parameters. Supplementary Fig. 4(a) shows the distribution of the 3 FWHMs of each bead image as a function of the position in the field of view.

The PSFs in the centre of the field of view (Supplementary Fig. 4(c)) are similar to Gaussian functions with FWHMs of around 650 nm in x and y and 3400 nm in z. The FWHMs significantly increase near the edge of the field of view with aberration, as shown in Supplementary Fig. 4(b,d).

### 5 Convolutional neural network for spectrum classification

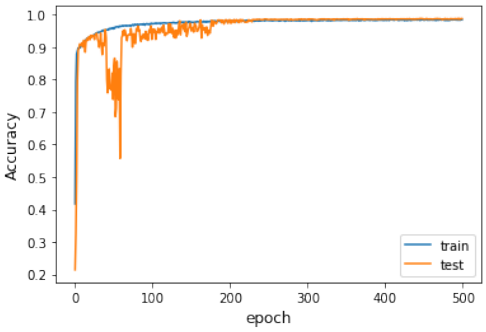


**Supplementary Figure 5** Training and validation accuracy of the CNN model.

#### 6 Fluorescence subtraction from spectra data

Autofluorescence signals from zebrafish embryos significantly reduces the contrast of Raman peaks. For better contrast, a broadband spectrum was estimated from each measured spectrum as the autofluorescence spectrum and subtracted.

The autofluorescence spectrum was estimated iteratively as follows: in each iteration, we first smoothed the data (MATLAB `smoothdata`) to obtain a broadband spectrum which was then compared with the original spectrum. For each point where the broadband spectrum was higher than the original spectrum, the intensity was lowered to the original value. The resulting spectrum was again smoothed. After 10 iterations, a broadband spectrum was obtained which approximately follows the valleys of the original spectrum. This spectrum was then used to estimate the autofluorescence and subtracted. Supplementary Fig. 6(a) shows a comparison between an original spectrum, estimated fluorescence signal and the final spectrum.

This method overestimates the fluorescence around 3000 $cm^{-1}$, particularly reducing the intensity of the water peak from the heart measurements. Therefore, for the heart measurements, an extra step was added to the iteration. Namely, the estimated fluorescence signal was flattened from 2760 $cm^{-1}$, as shown in Supplementary Fig. 6(b).

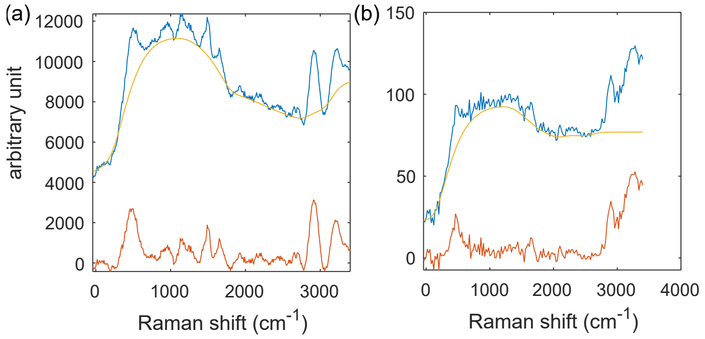


**Supplementary Figure 6 Fluorescence subtraction.** Example measured Raman spectra (blue) and the corresponding estimated fluorescence spectra (yellow) and the final spectra with fluorescence subtracted (red). (a) Spectra from the wound measurements. (b) Spectra from the heart measurement.

### 7 Comparison of heart images from the two cameras

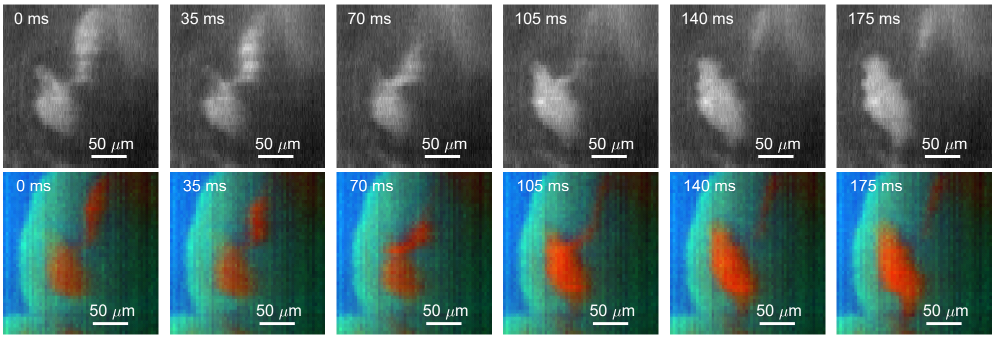


**Supplementary Figure 7 Comparison of heart images from the two cameras.** (Top): Raman images directly taken by Camera 1 at 6 time points, cropped and resized. (Bottom): corresponding processed Raman images obtained from hyperspectral images taken by Camera 2. There is a slight difference between the horizontal positions of the two image sets.
